# Optimization of an LNP-mRNA vaccine candidate targeting SARS-CoV-2 receptor-binding domain

**DOI:** 10.1101/2021.03.04.433852

**Authors:** Kouji Kobiyama, Masaki Imai, Nao Jounai, Misako Nakayama, Kou Hioki, Kiyoko Iwatsuki-Horimoto, Seiya Yamayoshi, Jun Tsuchida, Takako Niwa, Takashi Suzuki, Mutsumi Ito, Shinya Yamada, Tokiko Watanabe, Maki Kiso, Hideo Negishi, Burcu Temizoz, Hirohito Ishigaki, Yoshinori Kitagawa, Cong Thanh Nguyen, Yasushi Itoh, Fumihiko Takeshita, Yoshihiro Kawaoka, Ken J. Ishii

**Author notes:** Correspondence should be addressed to Ken J. Ishii, or Yoshihiro Kawaoka, Fumihiko Takeshita, Yasushi Itoh. The authors are contributed to this work equally (Kouji Kobiyama, Masaki Imai, Nao Jounai, Misako Nakayama).

## Abstract

In 2020, two mRNA-based vaccines, encoding the full length of severe acute respiratory syndrome coronavirus 2 (SARS-CoV-2) spike protein, have been introduced for control of the coronavirus disease (COVID-19) pandemic^1,2^. However, reactogenicity, such as fever, caused by innate immune responses to the vaccine formulation remains to be improved. Here, we optimized a lipid nanoparticle (LNP)-based mRNA vaccine candidate, encoding the SARS-CoV-2 spike protein receptor-binding domain (LNP-mRNA-RBD), which showed improved immunogenicity by removing reactogenic materials from the vaccine formulation and protective potential against SARS-CoV-2 infection in cynomolgus macaques. LNP-mRNA-RBD induced robust antigen-specific B cells and follicular helper T cells in the BALB/c strain but not in the C57BL/6 strain; the two strains have contrasting abilities to induce type I interferon production by dendritic cells. Removal of reactogenic materials from original synthesized mRNA by HPLC reduced type I interferon (IFN) production by dendritic cells, which improved immunogenicity. Immunization of cynomolgus macaques with an LNP encapsulating HPLC-purified mRNA induced robust anti-RBD IgG in the plasma and in various mucosal areas, including airways, thereby conferring protection against SARS-CoV-2 infection. Therefore, fine-tuning the balance between the immunogenic and reactogenic activity of mRNA-based vaccine formulations may offer safer and more efficacious outcomes.

---

The SARS-CoV-2 spike glycoprotein contains a receptor-binding domain (RBD) that binds to human angiotensin-converting enzyme 2 (hACE2) as a receptor to facilitate membrane fusion and cell entry^3^. Recent evidence suggests that the immune response to the SARS-CoV-2 spike protein is the key to controlling SARS-CoV-2 infection; a vaccine that can induce robust and specific T and B cells against the receptor-binding domain (RBD) of the spike protein antigen of SARS-CoV-2 may be ideal for protective efficacy and safety^4^. Accordingly, various animal experiments have demonstrated that the induction of humoral and cellular immune responses to the RBD by various types of vaccines confers protective efficacy with no signs of detrimental outcomes such as antibody-dependent enhancement^5,6^.

Concurrently with animal studies, a number of human clinical trials with various types of vaccines against COVID-19 have been initiated, conducted, and completed globally within a year after the viral genome sequence was reported in Wuhan, China, in December 2019^7^. Two mRNA vaccines encoding the full-length spike protein of SARS-CoV-2 have undergone Phase I-II-III trials, which were completed in nine months and approved by regulatory authorities in various countries as well as the WHO^1,2,8-10^. The results of their initial phase I–II clinical trials suggest that in both younger and older adults, the two vaccine candidates elicited similar dose-dependent SARS-CoV-2–neutralizing geometric mean titers, which were equivalent to that of a panel of SARS-CoV-2 convalescent serum samples^9,10^. It is worth noting that an mRNA vaccine (BNT162b2) encoding the full length of the SARS-CoV-2 spike protein was associated with a lower incidence and severity of systemic reactions than another mRNA vaccine encoding the RBD of spike protein (BNT162b1), particularly in older adults^10^. A few scientific explanations have been offered: one is the amount of mRNA in the RBD-mRNA vaccine, whose molar ratio is five times more than that of the full-length mRNA vaccine due to an RNA length shorter by 1/5 at the same dose. Although each RNA modification in the *in vitro* translated (IVT) mRNA to avoid innate immune recognition was made, the number or position of the modified nucleoside of the mRNAs may alter their immunostimulatory activity, acting as an endogenous adjuvant. Here, we optimized an mRNA vaccine candidate encoding SARS-CoV-2 spike protein RBD (319–541 aa) encapsulated in lipid nanoparticles (LNP-mRNA-RBD). To date, a mouse model using the BALB/c strain has been commonly used^6,11,12^, except for one study where C57BL/6 mice were immunized with LNP-mRNA encoding SARS-CoV-2 RBD, resulting in antigen-specific germinal center (GC) B cells and follicular helper CD4^+^ T cells (T_FH_) cells^13^. First, we immunized 6–8-week-old female mice of either the C57BL/6 or BALB/c strains intramuscularly with 3 μg of LNP-mRNA-RBD on days 0 and 14. Unexpectedly, after two intramuscular immunizations, LNP-mRNA-RBD induced significantly higher anti-RBD antibody responses in BALB/c mice but not in C57BL/6 mice in this study (**Fig. 1a, Extended Fig 1**). To understand why LNP-mRNA-RBD immunogenicity for antigen-specific B cell responses was different among mouse strains, we further examined whether LNP-mRNA-RBD induces T_FH_ and GC B cells collected from the draining popliteal lymph nodes (pLN) and analyzed by flow cytometry (**Extended data Fig. 2)**. In correlation with serum antibody responses, the frequency (%) of both T_FH_ (CD4^+^CD185^+^PD-1^+^ cells) and GC B cells (CD38^-^GL7^+^CD19^+^ cells) in the immunized pLN was significantly higher in BALB/c mice than that in C57BL/6 mice (**Fig. 1b-e**) after LNP-mRNA-RBD immunization. To further evaluate antigen-specific CD8^+^ and other CD4^+^ T cells induced by LNP-mRNA-RBD, we synthesized 128 peptides, consisting of a 20-aa sequence of spike protein with 10 overlapping aa divided into eight pools containing 16 peptides in one pool (**Fig. 1f**). After two LNP-mRNA-RBD immunizations in either mouse strain, *in vitro* re-stimulation of the immunized spleen with peptide pools 3 and 4 induced substantial IFN-γ production in C57BL/6 mice, while in BALB/c mice this was achieved with peptide pool 3 (**Fig. 1g and h, Extended data Fig. 3a and b**). IL-13 production was not found in the supernatant of the spleen cell culture with peptides in either C57BL/6 or BALB/c mice (**Extended data Fig. 3c and d**). To further characterize LNP-mRNA-RBD-induced T cells, we performed intracellular cytokine staining of three antigen-specific type-1 cytokines (IL-2, IFN-γ, and TNF-α) produced by the immunized spleen T cells re-stimulated with peptide pools 2, 3, or 4. Spike antigen-specific polyfunctional CD8^+^ and CD4^+^ T cells were significantly upregulated in LNP-mRNA-RBD-immunized BALB/c mice after re-stimulating the spleen cells with peptide pools 3 and 4 (**Fig. 1h, Extended data Fig. 4b and 5b**). However, those in the immunized C57BL/6 mice showed substantial polyfunctional CD8^+^ T cells and weak CD4^+^ T cell responses (**Fig. 1g, Extended data Fig. 4a and 5a**). These data clearly demonstrate that LNP-mRNA-RBD induces robust B and T cell responses in BALB/c mice but relatively weak T cell and B cell responses in C57BL/6 mice, suggesting the immunogenic profile of LNP-mRNA-RBD is different between these mouse strains.

**Figure 1.**
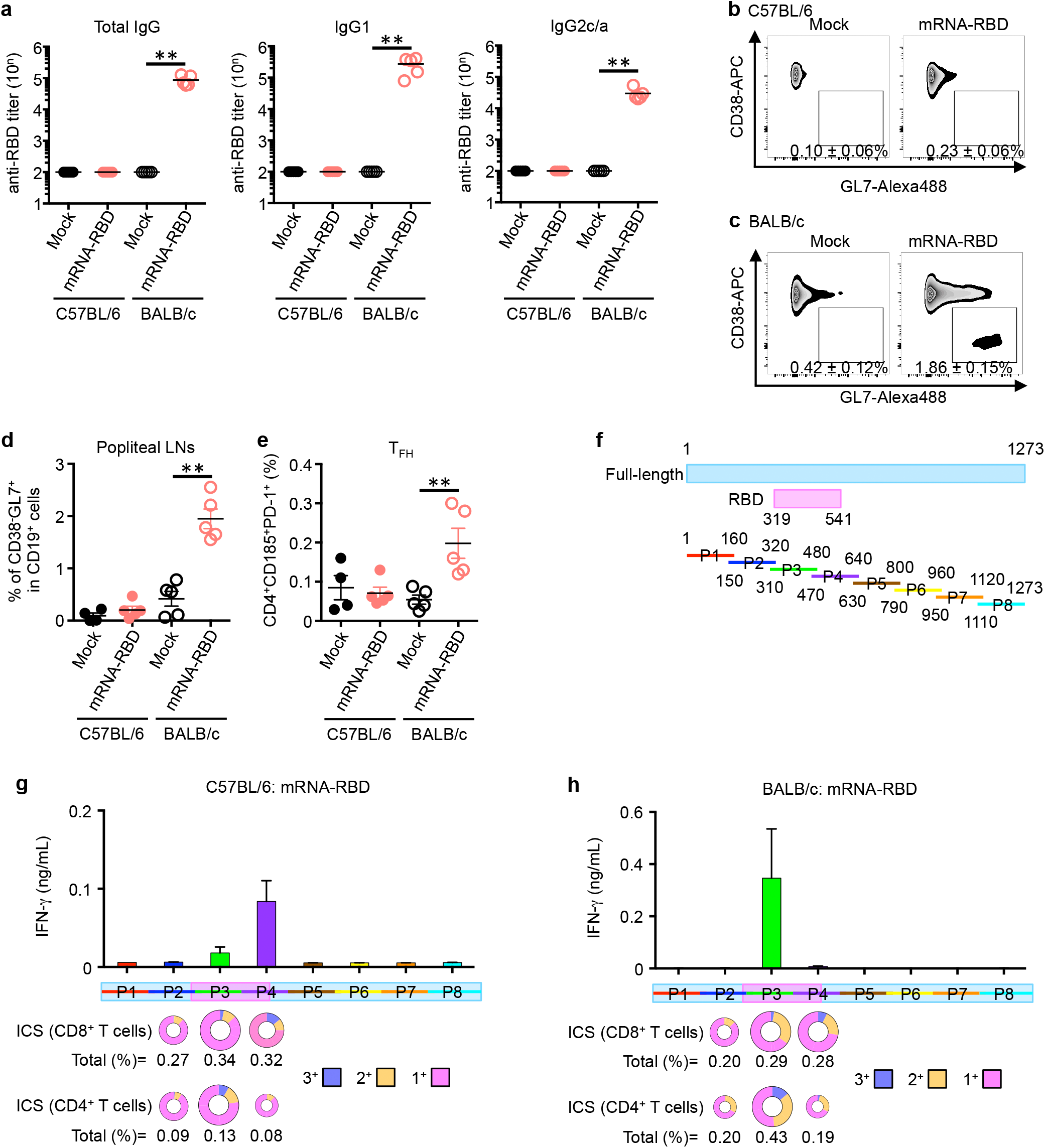
Mouse strain-specific immunogenicity of mRNA vaccine against SARS-CoV-2 RBD. (**a–e, g, and h**) Six to eight week-old C57BL/6 and BALB/c mice were intramuscularly immunized with mock or LNP-mRNA-RBD (3 μg) at days 0 and 14. (**a)** Two weeks after the second immunization, plasma anti-RBD antibody titers were measured using ELISA. (**b**–**e**) Popliteal lymph nodes were collected from immunized mice. (**b**–**d**) GC B cells were gated as GL7^+^CD38^-^CD19^+^ cells. (**e**) T_FH_ cells were gated as CD185^+^PD-1^+^CD3ε^+^CD4^+^ T cells. (**f**) Overlapping peptides of SARS-CoV-2 spike protein. Overlapping peptides were divided into eight pools, and each pool contained 16 peptides. (**g**–**h**) Cells were harvested from the spleen of mRNA-RBD immunized mice and re-stimulated with pooled peptides for 24 h. IFN-γ levels in the culture supernatant were measured using ELISA. (**g**–**h**) Percentages of cytokine-producing CD8^+^ and CD4^+^ T cells after stimulation with pools 2, 3, and 4 for 6 h with protein transport inhibitor are shown in a pie chart. 3^+^: IFN-γ^+^IL-2^+^TNF-α^+^, 2^+^: IFN-γ^+^IL-2^+^, IFN-γ^+^TNF-α^+^, and IL-2^+^TNF-α^+^, 1^+^: IFN-γ^+^, IL-2^+^, and TNF-α^+^. *N* = 4–5 mice per group, mean ± SEM, \**p* < 0.05 by Mann-Whitney test.

Nucleic acid-based vaccines are known to utilize their backbone DNA or RNA as built-in adjuvants^14-16^. In LNP-mRNA vaccines, it has been shown that mRNA itself acts as an endogenous adjuvant sensed by Toll-like receptors 3, 7, or 8 and/or cytosolic RNA sensors such as RIG-I and MDA5^17^. Kariko et al. reported that modification of RNA by methylation or incorporating modified nucleoside such as pseudouridine enables the escape from innate immune sensing, thereby improving translation efficiency^18,19^. Several studies have revealed that type I IFN interferes with the CD8 T cell responses elicited by LNP-mRNA and the translation efficiency of the encoded protein^20,21,22^. In addition to T cell responses, BNT162b1 showed higher reactogenicity than BNT162b2 in the clinical trial; therefore, BNT162b2 has been selected for further development in a Phase III clinical trial^10^. The reason for the difference in reactogenicity remains unclear, but the authors considered that immunostimulatory activity of the mRNA in LNP formulation might be attributed to its reactogenicity^10^.

In order to translate our findings from mice to humans, we then examined whether LNP-mRNA-RBD triggers type I IFN production from human PBMCs. When mixed with LNP-mRNA-RBD *in vitro*, PBMCs from three healthy humans produced a higher amount of IFN-α than that induced by LNP-mRNA-Full (**Fig. 2a**). We then performed a similar experiment using mouse bone marrow-derived dendritic cells (BM-DCs) from either C57BL/6 or BALB/c mice. Surprisingly, a high level of IFN-α was observed upon culture with LNP-mRNA-full or LNP-mRNA-RBD in C57BL/6 mice, but very low or no IFN-α production was observed in BALB/c mice (**Fig. 2b**). LNP-mRNA products usually contain undesirable RNA, such as dsRNA as TLR3 ligand^22^, produced during the manufacturing process, which might affect innate immune activation. To remove such RNA byproducts, we performed HPLC purification (**data not shown**) and then the resultant mRNA containing the active ingredient was encapsulated in LNP [RBD (HPLC)]. RBD (HPLC) showed significantly less potential in production of type I IFN from both human PBMCs and GM-DCs (**Fig. 2a and b**). In order to examine the immunogenicity, C57BL/6 or BALB/c mice were administered with RBD (HPLC) or LNP-mRNA-RBD. Of interest, RBD (HPLC) showed significantly higher levels of the RBD-specific B cell response than LNP-mRNA-RBD, including serum IgG1, IgG2, and total IgG in both BALB/c and C57BL/6 mice (**Fig. 2c and Extended Fig. 6a**). In particular, RBD (HPLC) induced significantly higher number of GC B cells in the draining lymph nodes of the C57BL/6 mice than LNP-mRNA-RBD (**Fig. 2d and e**). In addition to antibody responses, effects of RNA-purification on antigen-specific T cell responses were further examined. RBD (HPLC) induced higher frequency of the RBD-specific polyfunctional CD8^+^ and CD4^+^ T cells that produced significantly more IFN-γ and other type-1 cytokines, but not type-2 cytokines such as IL-13, in response to peptide pools 3 or 4 re-stimulation than LNP-mRNA-RBD (**Fig. 2f-i, and Extended Fig. 6b–e, 7, 8**).

**Figure 2.**
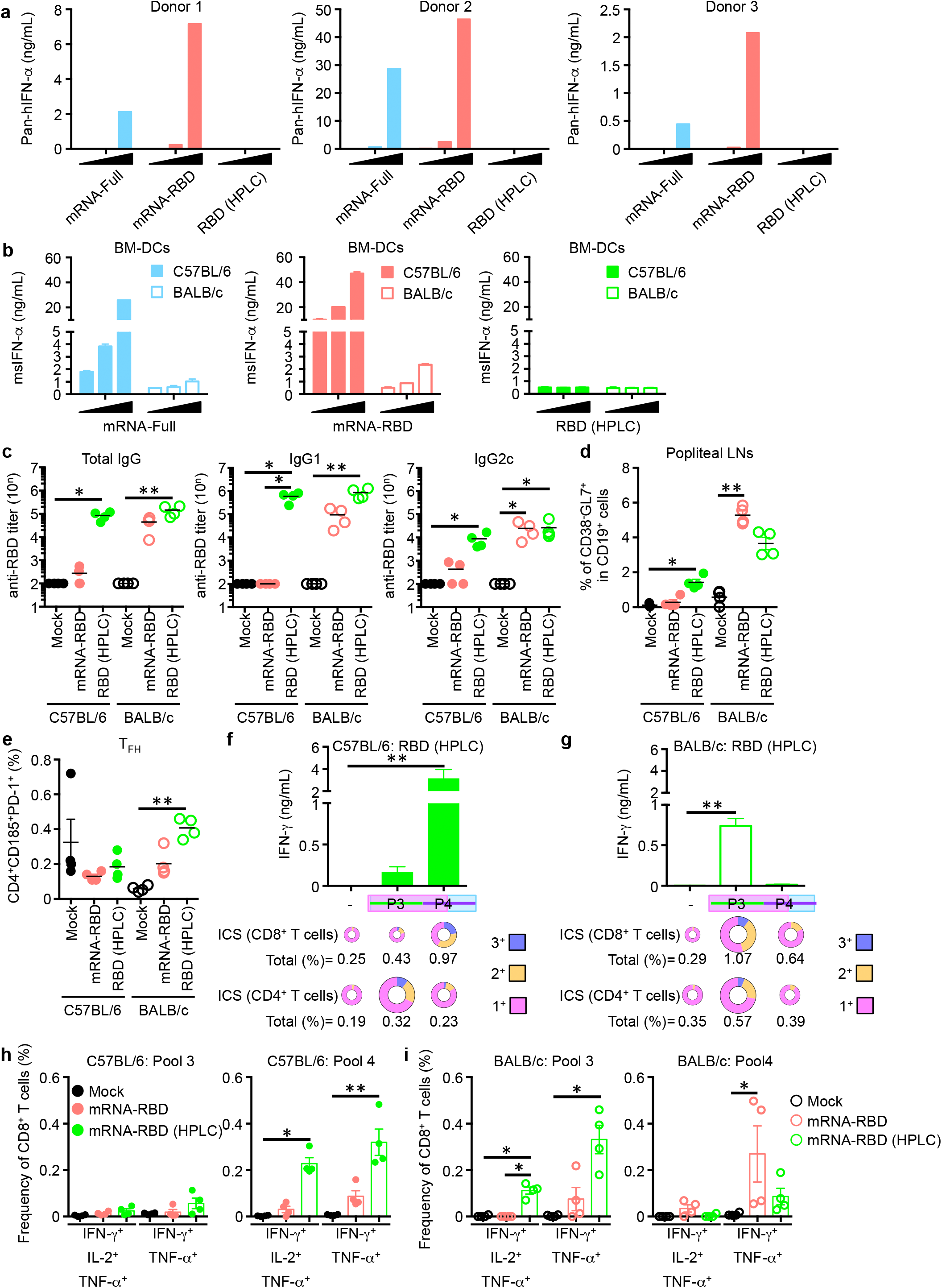
HPLC purification improves the immunogenicity of mRNA vaccine. (**a**) Human PBMCs from non-infected individuals were stimulated with LNP-mRNA-Full (0.4, 2, and 10 μg/mL), LNP-mRNA-RBD (0.4, 2, and 10 μg/mL), or LNP-mRNA-RBD (HPLC) (0.4, 2, and 10 μg/mL) for 24 h. IFN-α level in the culture supernatant was measured using ELISA. (**b**) Bone-marrow-derived dendritic cells (BM-DCs) from C57BL/6 and BALB/c mice were stimulated by LNP-mRNA-Full (0.4, 2, and 10 μg/mL), LNP-mRNA-RBD (0.4, 2, and 10 μg/mL), or LNP-mRNA-RBD (HPLC) (0.4, 2, and 10 μg/mL) for 24 h. IFN-α level in the culture supernatant was measured using ELISA. (**c–i**) C57BL/6 mice were intramuscularly immunized with mock, LNP-mRNA-RBD (3 μg), or LNP-mRNA-RBD (HPLC) (3 μg) at days 0 and 14. (**c**) Two weeks after the second immunization, plasma anti-RBD antibody titers were measured using ELISA. (**d** and **e**) Popliteal lymph nodes were collected from immunized mice. (**d)** GC B cells were gated as GL7^+^CD38^-^CD19^+^ cells. (**e**) T_FH_ cells were gated as CD185^+^PD-1^+^CD3ε^+^CD4^+^ T cells. (**f** and **g**) Cells were harvested from the spleen of immunized mice and re-stimulated with pooled peptides for 24 h. IFN-γ level in the culture supernatant was measured using ELISA. Percentages of cytokine-producing CD8^+^ and CD4^+^ T cells after stimulation of peptide pools 3 and 4 for 6 h with protein transport inhibitors are shown in a pie chart. 3^+^: IFN-γ^+^IL-2^+^TNF-α^+^, 2^+^: IFN-γ^+^IL-2^+^, IFN-γ^+^TNF-α^+^, and IL-2^+^TNF-α^+^, 1^+^: IFN-γ^+^, IL-2^+^, and TNF-α^+^. (**h** and **i**) Representative data from Figure 2f, g, Extended data Fig. 8, and 9 are shown. IFN-γ^+^IL-2^+^TNF-α^+^ and IFN-γ^+^TNF-α^+^ CD8^+^ T cell are shown as a scatter dot plot. *N* = 4–5 mice per group, mean ± SEM, * *p* <0 .05 by ANOVA followed by Dunn’s multiple comparisons test.

To further translate these findings to a more relevant pre-clinical evaluation of RBD (HPLC), non-human primates (NHPs), cynomolgus macaques, were chosen for further study. In this study, we immunized four macaques intramuscularly with RBD (HPLC) with two macaques as mock controls. After the first immunization, RBD (HPLC) induced an anti-RBD-specific antibody, and the second immunization enhanced these responses (**Fig. 3b**). Neutralizing antibodies were also induced by RBD (HPLC) vaccination (**Fig. 3c**). We further examined antigen-specific antibody responses in swab samples. Interestingly, following intramuscular immunization, levels of RBD-specific IgG in the swab samples from conjunctiva, nasal cavity, oral cavity, trachea, and rectum were significantly higher in RBD (HPLC) group than in the mock group (**Fig. 3d**). Individual macaques administered with RBD (HPLC) showed drastically lower RNA levels of SARS-CoV-2 (**Fig. 4a**) and infectious virus (**Fig. 4b**) in the swab at day 1 post-infection than those administered with mock. Viral RNA levels in the trachea, bronchus, and lung were lower in vaccinated macaques at day 7 (**Fig. 4c and Extended Fig. 9**). All mock-administered macaques manifested fever and pneumonia after viral infection, which were not observed in immunized macaques (**Extended Fig. 10 and 11**). These results suggest that RBD (HPLC) administration confers protection against SARS-CoV-2 infection. Histological analysis of the lung at 7 days post infection demonstrated infiltration of lymphocytes and neutrophils, alveolar wall thickening, and viral protein in macaques administered with mock but not in those administered with RBD (HPLC) (**Fig. 4d and 4e**). Accordingly, histological scores of the lung in macaques administered with RBD (HPLC) were lower than those administered with mock (**Fig. 4f**). Of importance, intramuscular administration with RBD (HPLC) induced the development of bronchus-associated lymphoid tissue (BALT) (**Fig. 4d**), although intramuscular immunization induced RBD-specific IgG, but not IgA, in swab samples without intranasal and intratracheal virus challenge (**Fig. 3b**). Interestingly, the IgG titer was slightly reduced or similar after viral challenge (**Fig. 3d**). These results suggest that the induced antibody in the mucosa through BALT formation, such as the nasal and trachea mucosa, might capture and neutralize SARS-CoV-2, resulting in the reduction of viral RNA and infective virus in the swab at day 1 post-challenge.

**Figure 3.**
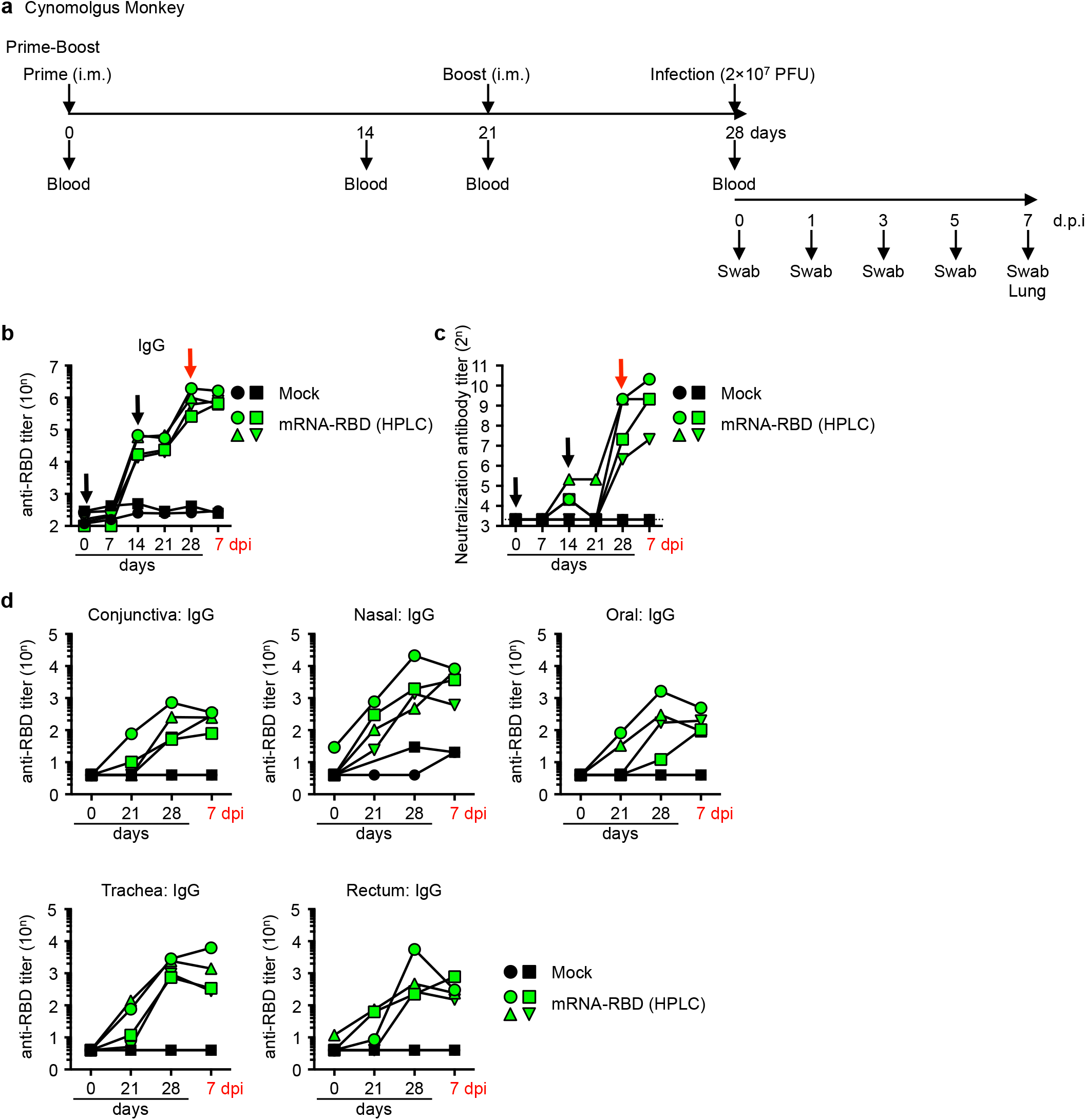
HPLC-purified LNP-mRNA-RBD induces RBD-specific antibodies in the plasma and swab samples of non-human primates. (**a**) Schedule of immunization, infection, and sample collection. (**b**–**c**) Cynomolgus macaques were intramuscularly immunized with mock or LNP-mRNA-RBD (HPLC) (100 μg) at days 0 and 21. (**b**) Plasma anti-RBD antibody titer at days 0, 7, 14, 21, 28, and 7 dpi were measured using ELISA. (**c**) Neutralizing activity against SARS-CoV-2 infection were measured by neutralization assay. (**d**) Anti-RBD IgG titers in the swab samples (conjunctiva, oral cavity, nasal cavity, tracheal, and rectum) were measured using ELISA. Black arrows indicate date of vaccination, and red arrows indicate infection date.

**Figure 4.**
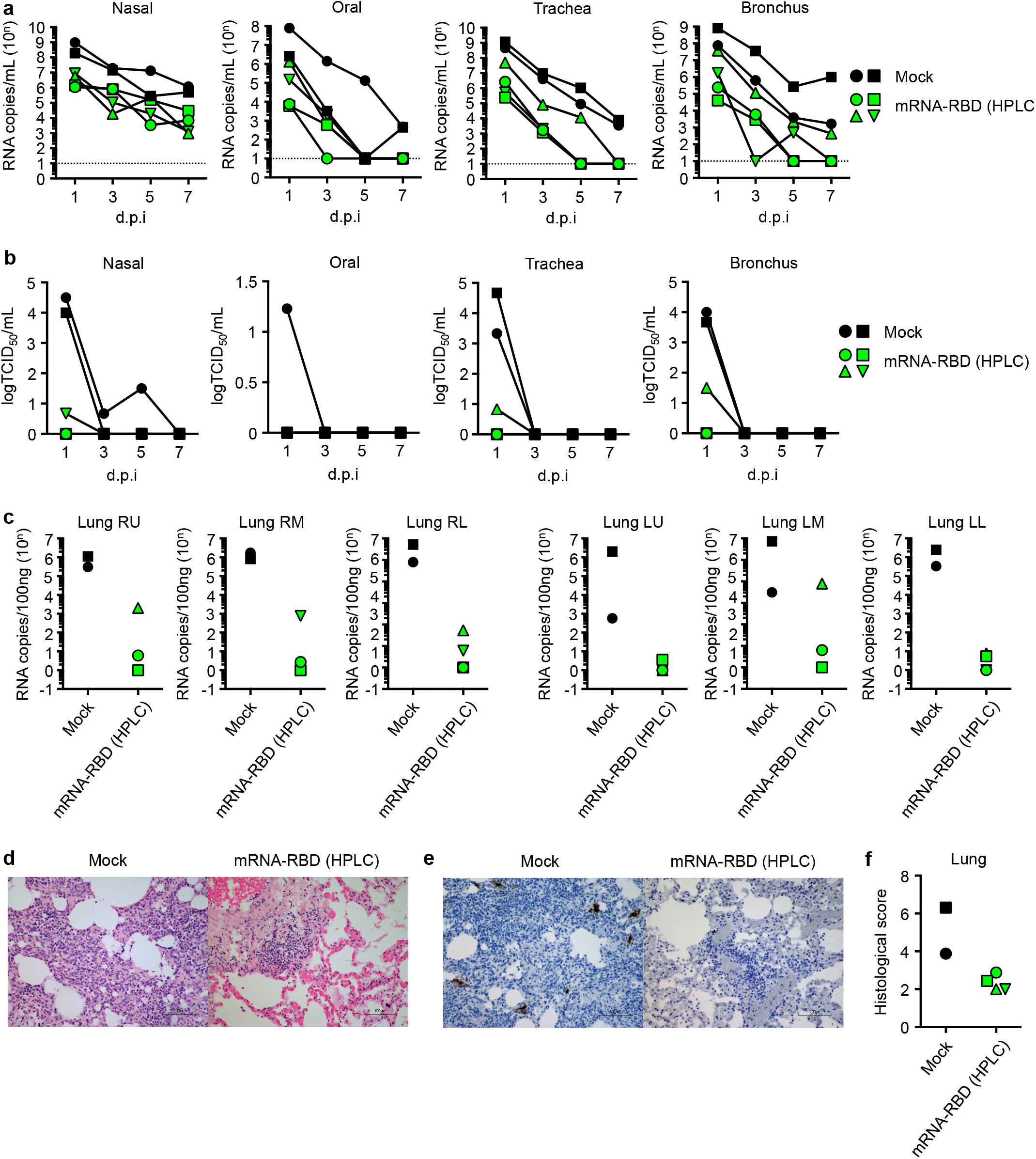
HPLC-purified LNP-mRNA-RBD protects against SARS-CoV-2 infection in non-human primates. One week after the second immunization, SARS-CoV-2 (2 × 10^7^ PFU) was inoculated into conjunctiva, nasal cavity, oral cavity, and trachea of cynomolgus. (**a**) Viral RNA and (**b**) viral titers in the swab sample were measured by RT-PCR and a cell culture method. (**c**) Viral RNA in the lung tissues were measured by RT-PCR. RU: right upper lobe, RM: right middle lobe, RL: right lower lobe, LU: left upper lobe, LM: left middle lobe, LL: left lower lobe. (**d**) HE staining and (**e**) immunohistochemical staining of viral nucleocapsid protein in lung sections from Mock (left) and mRNA-RBD (HPLC) (right) immunized macaques. (**f**) The average histological scores of eight sections in each macaque were evaluated in a blinded manner.

In this study, we evaluated the nonclinical efficacy of LNP-mRNA vaccine candidates targeting SARS-CoV-2 RBD. First, LNP-mRNA-RBD showed higher immunogenicity only in BALB/c mice than in C57BL/6 mice (**Fig. 1a**). We initially interpreted the data by suggesting the less T cell epitopes of the RBD in C57/BL6 as the cause. In fact, CD4 Tfh induction was lower in C57BL/6 mice than that of BALB/c mice even after HPLC purification (**Fig.1e and 2e**). However, recent clinical trials by BioNTech/Pfizer showed that an mRNA vaccine that encoded the RBD resulted in a high titer of RBD-specific IgG and neutralizing antibodies in humans and monkeys. These results suggest that RBD does contain T cell epitopes, at least in primates^6,10^. These data led us to hypothesize that the difference in vaccine-induced adaptive immune responses is altered by the species- or strain-specific innate immune responses to the LNP-mRNA formulation, which is shown to interfere with the mRNA expression of the protein antigen of interest, thereby reducing immunogenicity and efficacy^16^. Our data strongly suggest that optimization of purification and formulation of LNP-mRNA contributes to improvement of LNP-mRNA immunogenicity with less reactogenicity. It is of note that macaques administered with LNP-mRNA targeting RBD acquired significantly high levels of protective IgG specific to SARS-CoV-2 in mucosal swab samples from conjunctiva, oral cavity, nasal cavity, trachea, bronchus, and rectum (**Fig. 3d** and data not shown). Corbett KS et al. recently demonstrated that vaccination of NHPs with LNP-mRNA encoding the full-length spike antigen (mRNA-1273) induced robust SARS-CoV-2 neutralizing activity and rapid protection in the upper and lower airways and showed that the IgG level in the BALF was higher than the IgA level after the infection, although whether the vaccine antigen-specific IgG was induced before the virus challenge was not shown^23^. Although HPLC-purified LNP-mRNA-RBD elicited RBD-specific mucosal IgG, no RBD-specific IgA was detected (data not shown), indicating that the mucosal IgG through BALT formation or leaked from the blood circulation, which may be critical for the protective efficacy of LNP-mRNA-RBD. Further detailed analyses are needed to clarify whether LNP-mRNA induces unique and/or specific immune responses including IgG secretion in the mucosa after intramuscular vaccination.

Based on our results obtained in murine and NHP models, reduction of reactogenicity without losing immunogenicity, in other words, fine-tuning of the balance between endogenous adjuvant activity and antigen translation efficiency of LNP-mRNA, may provide a means towards better efficacy and safety and will also be crucial for the development of anti-SARS-CoV2 vaccines in the near future.

## Figure legends

**Extended data Fig 1.**
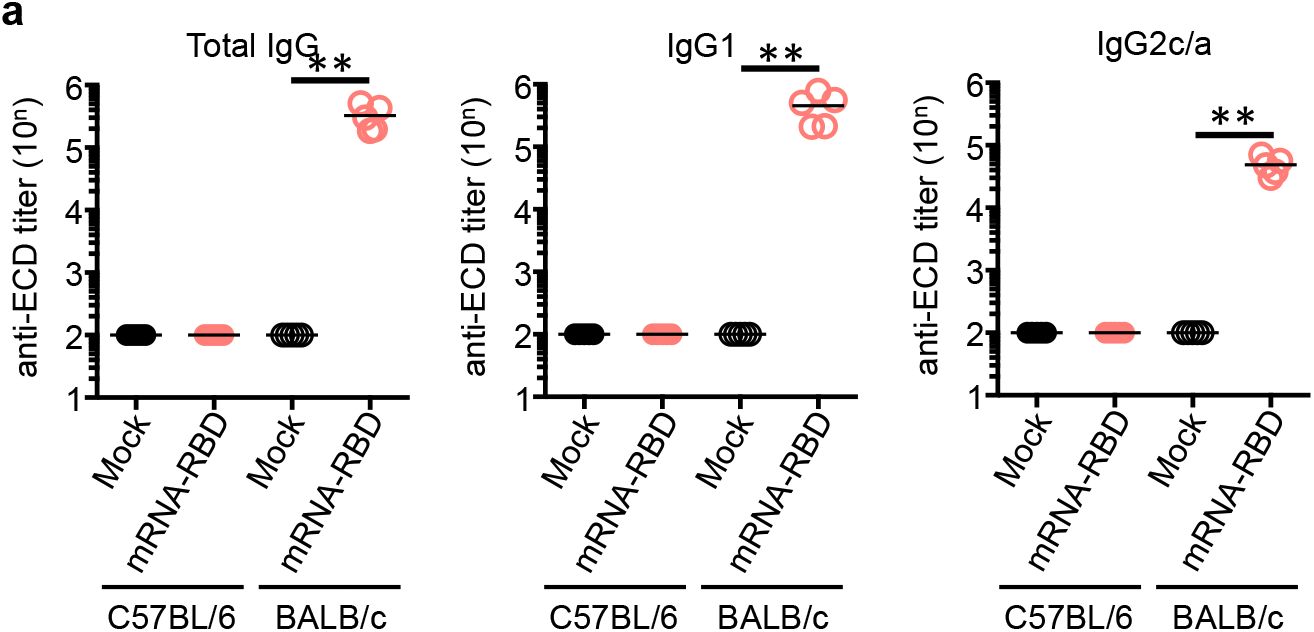
LNP-mRNA-RBD vaccine induces ectodomain-specific antibody responses in BALB/c mice. C57BL/6 and BALB/c mice were intramuscularly immunized with mock or LNP-mRNA-RBD (3 μg) on days 0 and 14. Two weeks after the second immunization, plasma anti-ECD antibody titers were measured using ELISA. *N* = 4–5 mice per group, mean ± SEM, \**p* < 0.05 by Mann-Whitney test.

**Extended data Fig 2.**
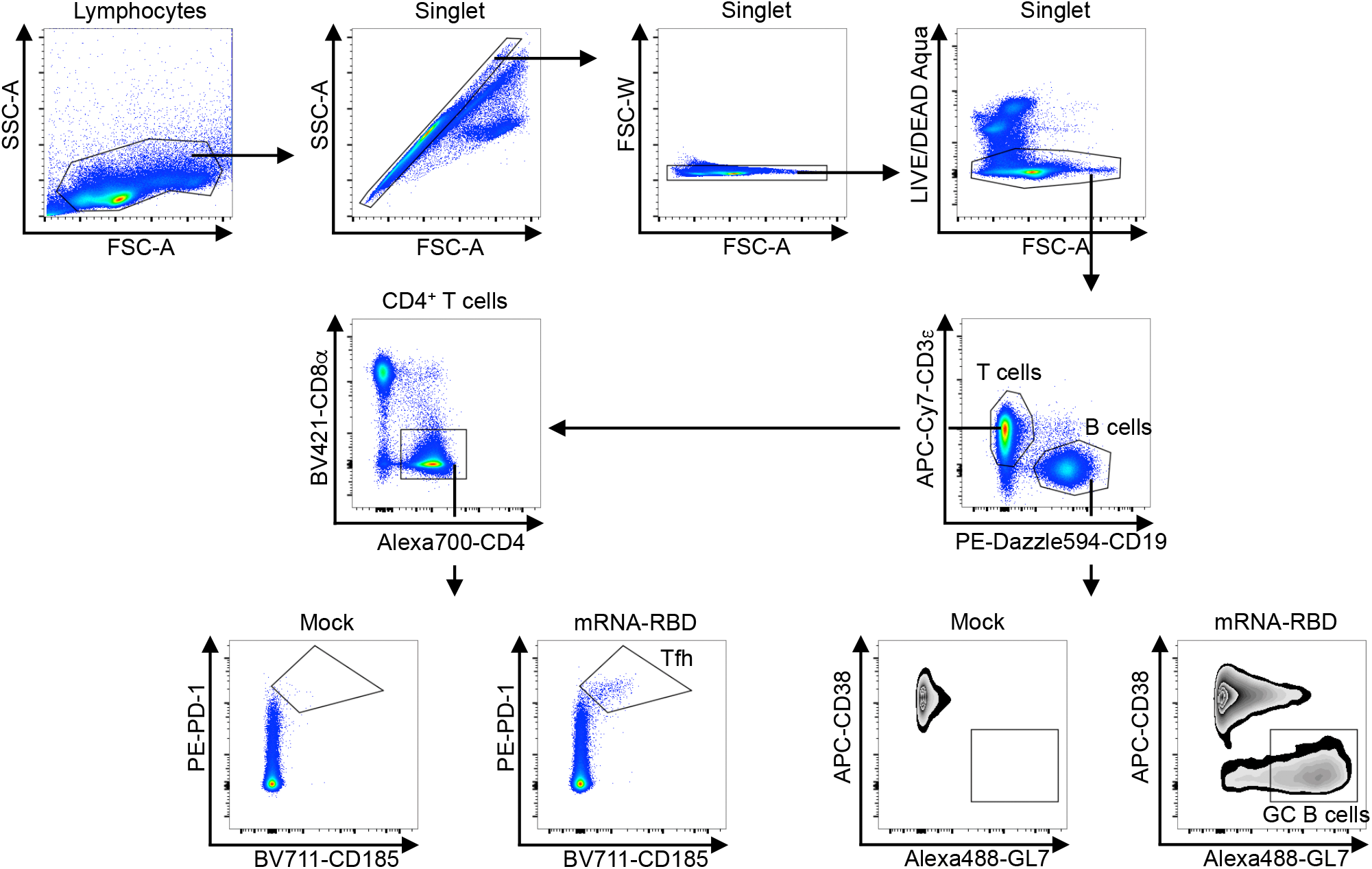
Gating strategy for GC B and T_FH_ cells. Cells were harvested from popliteal lymph nodes of immunized mice and stained for GC B and T_FH_ cells. Cells were gated for lymphocyte size, singlets, live, T or B cells, and T_FH_ or GC B cells.

**Extended data Fig 3.**
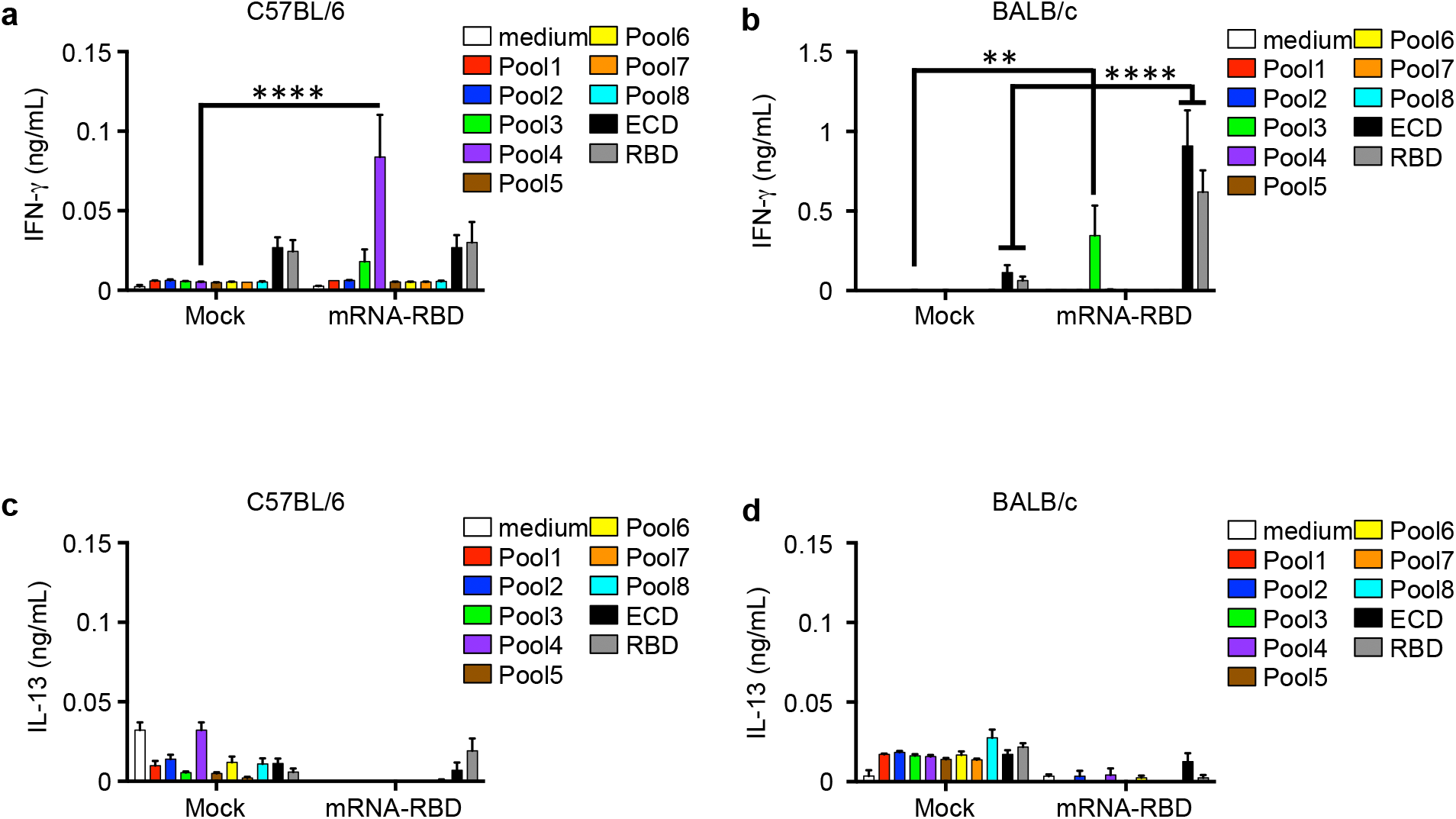
T cell responses to LNP-mRNA-RBD. Cells were harvested from the spleen of mRNA-immunized mice, re-stimulated by the spike protein peptide pool, ECD, or RBD for 24 h. IFN-γ and IL-13 levels in the culture supernatant were measured using ELISA. *N* = 4–5 mice per group, mean ± SEM, * *p* < 0.05 by ANOVA followed by Sidak’s multiple comparisons test.

**Extended data Fig 4.**
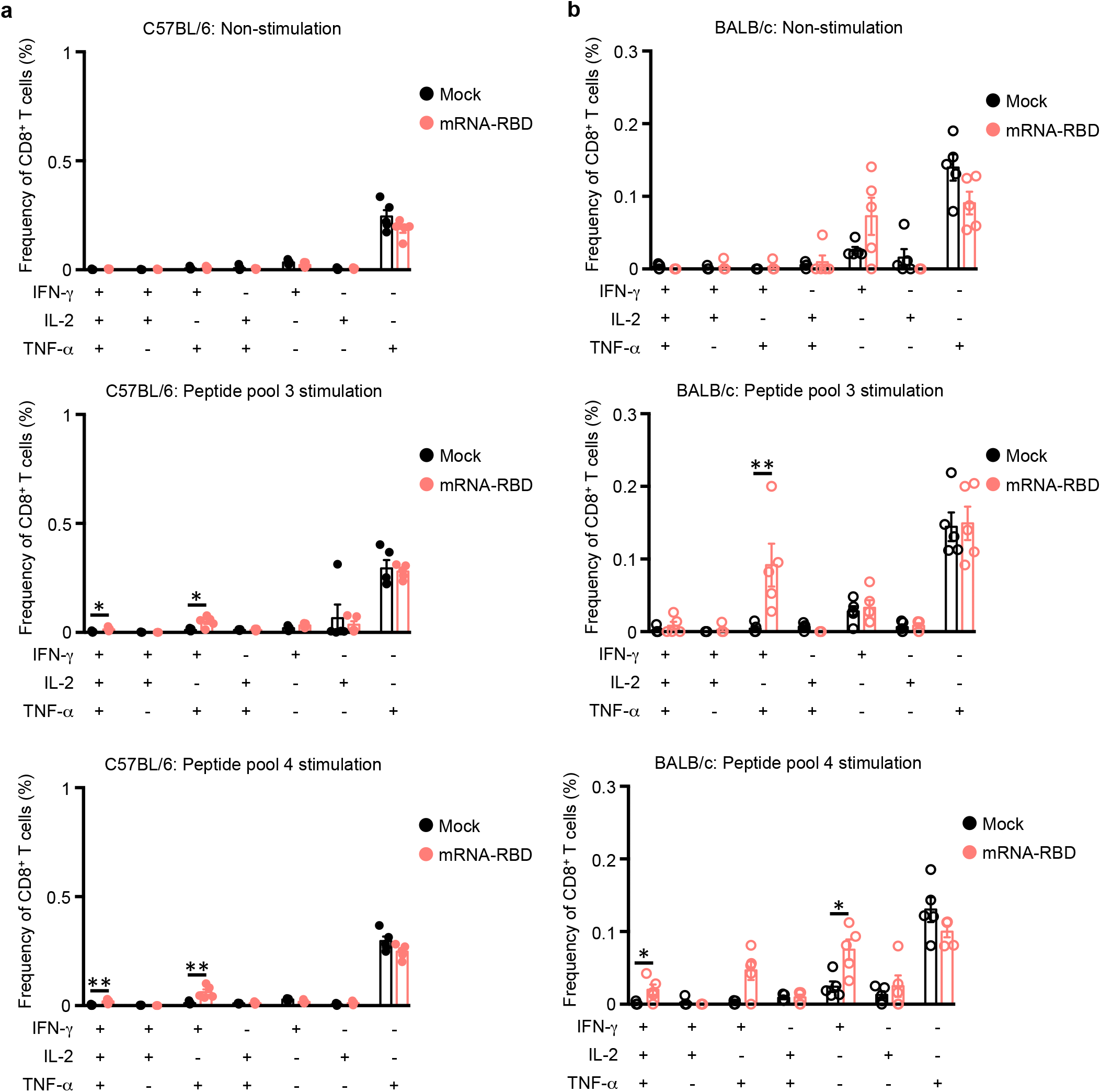
CD8 T cell responses to the mRNA vaccine. Cells were harvested from the spleen of immunized mice and re-stimulated by pooled peptides for 6 h with a protein transport inhibitor. The percentage of cytokine-producing CD8^+^ T cells was analyzed by flow cytometry. *N* = 4–5 mice per group, mean ± SEM, \**p* < 0.05 by Mann-Whitney test.

**Extended data Fig 5.**
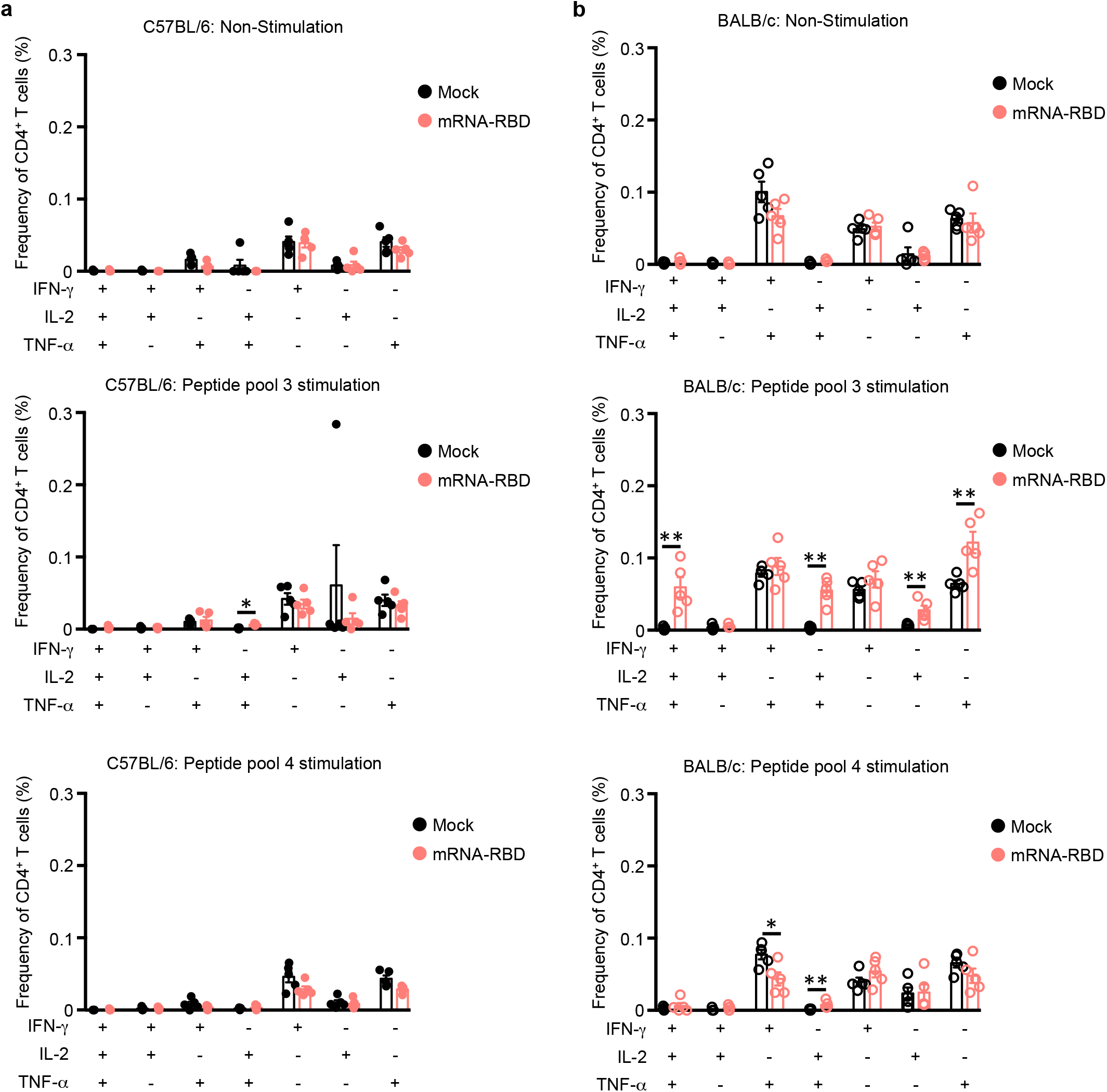
CD4 T cell responses to the mRNA vaccine. Cells were harvested from the spleen of immunized mice and re-stimulated by pooled peptides for 6 h with a protein transport inhibitor. Percentage of cytokine-producing CD4^+^ T cells was analyzed by flow cytometry. *N* = 4–5 mice per group, mean ± SEM, \**p* < 0.05 by Mann-Whitney test.

**Extended data Fig 6.**
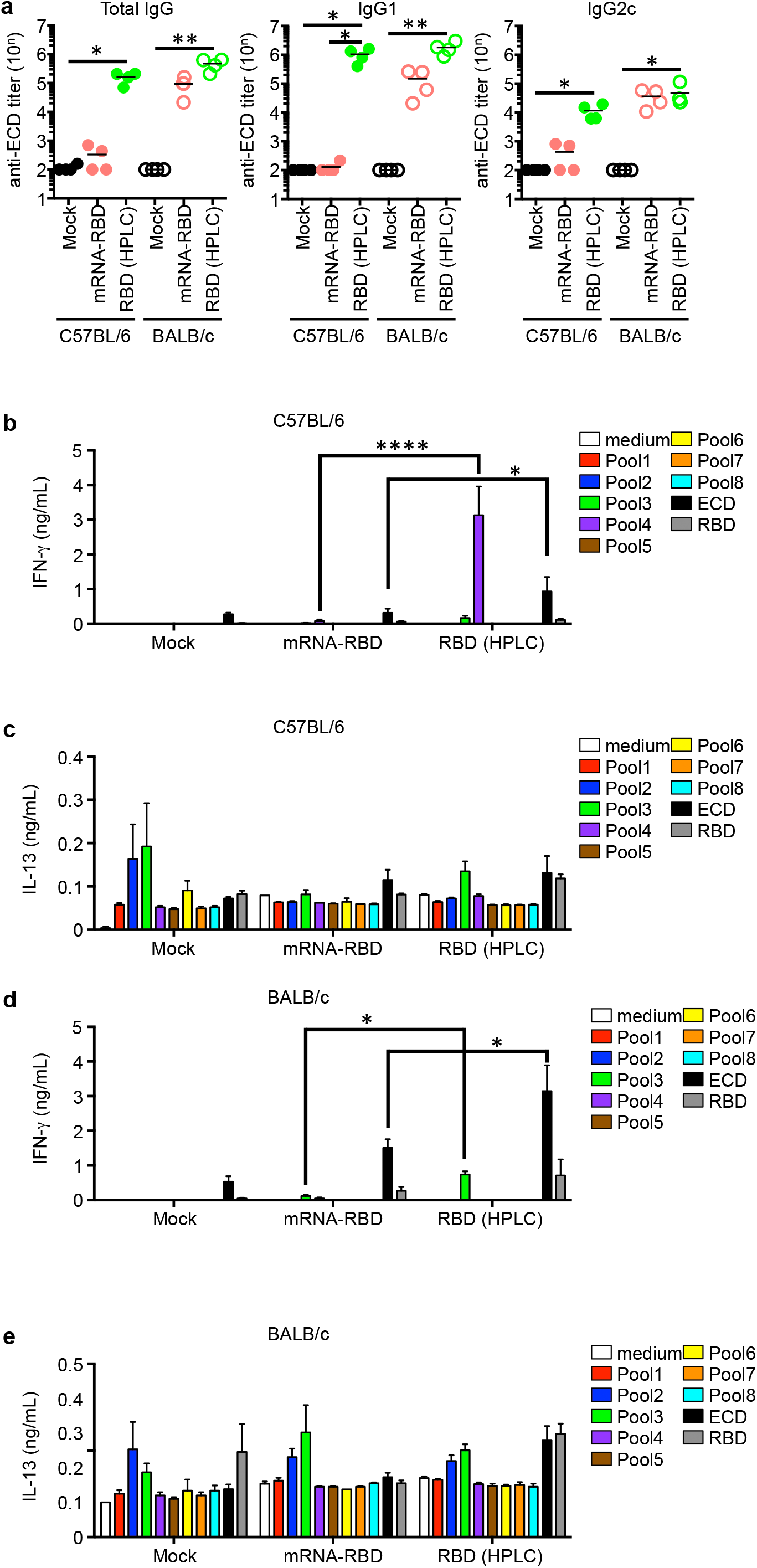
T cell responses to an HPLC-purified mRNA vaccine in C57/BL6 mice. (**a**) C57/BL6 and BALB/c mice were intramuscularly immunized with mock, mRNA-RBD, or RBD (HPLC) (3 μg) on days 0 and 14. Two weeks after the second immunization, serum anti-ECD antibody titers were measured using ELISA. (**b– e**) Cells were harvested from the spleen of mRNA-immunized mice, re-stimulated by the peptide pool of spike protein, ECD, or RBD for 24 h. IFN-γ and IL-13 levels in the culture supernatant were measured using ELISA. *N* = 4 mice per group, mean ± SEM, * *p* < 0.05 by ANOVA followed by Dunn’s or Sidak’s multiple comparisons test.

**Extended data Fig 7.**
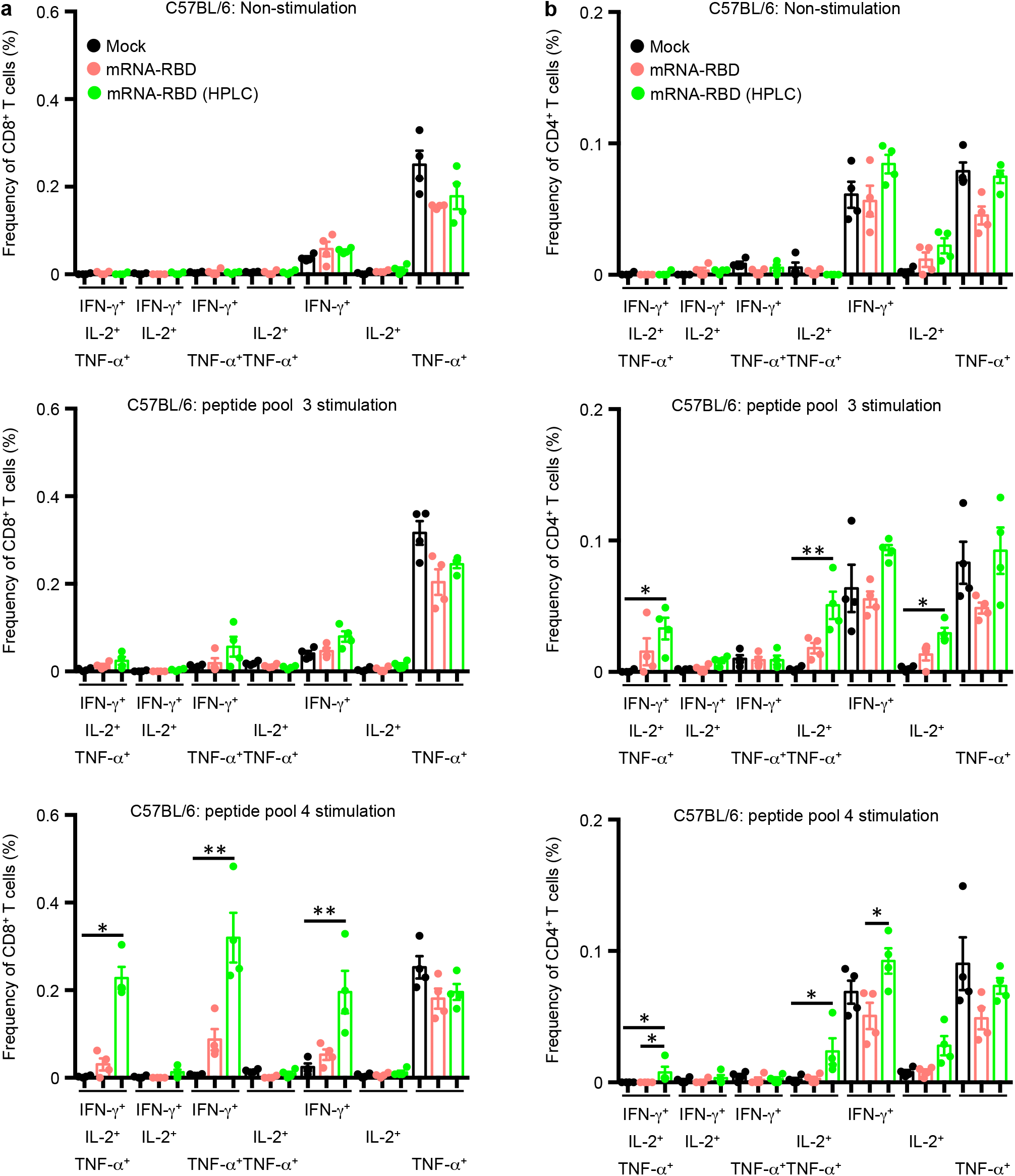
T cell responses to an HPLC-purified mRNA vaccine in C57/BL6 mice. Cells were harvested from the spleen of immunized mice and re-stimulated by pooled peptides for 6 h with a protein transport inhibitor. The percentage of cytokine-producing CD8^+^ and CD4^+^ T cells was analyzed by flow cytometry. *N* = 4 mice per group, mean ± SEM, * *p* < 0.05 by ANOVA followed by Dunn’s multiple comparisons test.

**Extended data Fig 8.**
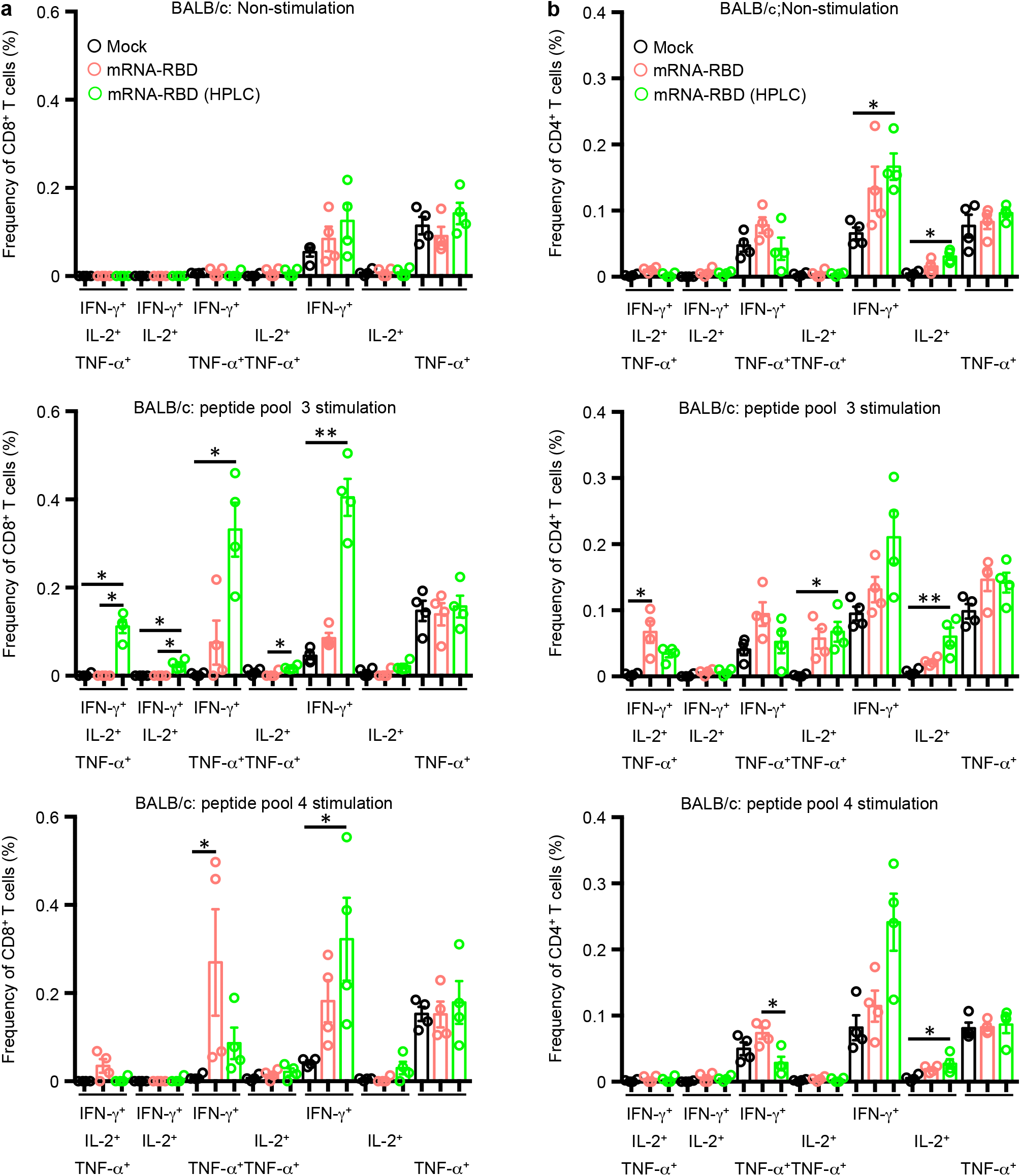
T cell responses to an HPLC-purified mRNA vaccine in BALB/c mice. Cells were harvested from the spleen of immunized mice and re-stimulated with pooled peptides for 6 h with a protein transport inhibitor. The percentage of cytokine-producing CD8^+^ and CD4^+^ T cells was analyzed by flow cytometry. *N* = 4 mice per group, mean ± SEM, * *p* < 0.05 by ANOVA followed by Dunn’s multiple comparisons test.

**Extended data Fig 9.**
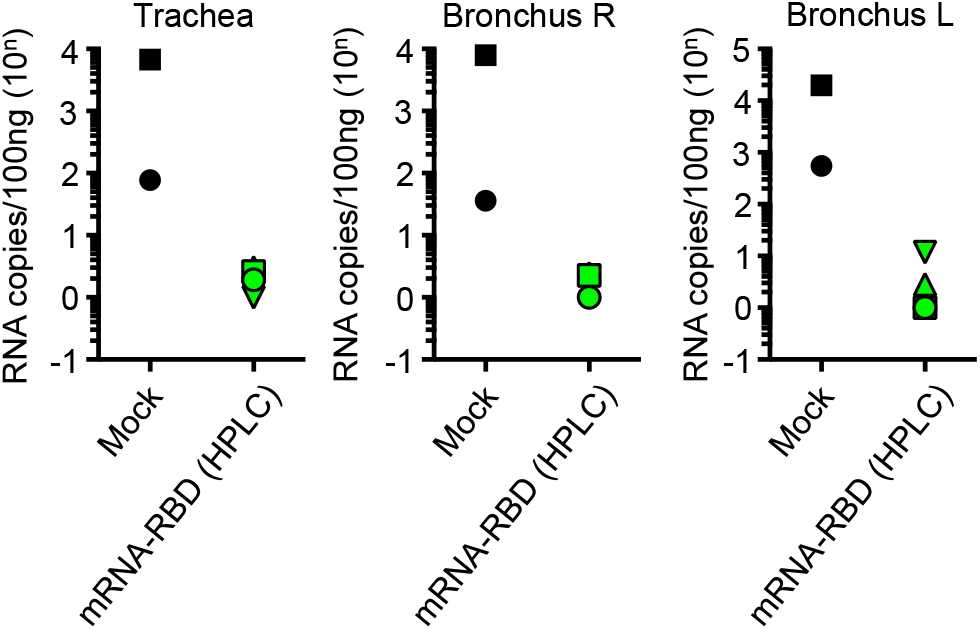
HPLC-purified mRNA vaccine protects against SARS-CoV-2 infection in non-human primates. One week after the second immunization, SARS-CoV-2 (2 × 10^7^ PFU) was inoculated into conjunctiva, nasal cavity, oral cavity, and trachea of cynomolgus. Viral RNA in the trachea and bronchus tissues were measured by RT-PCR.

**Extended data Fig 10.**
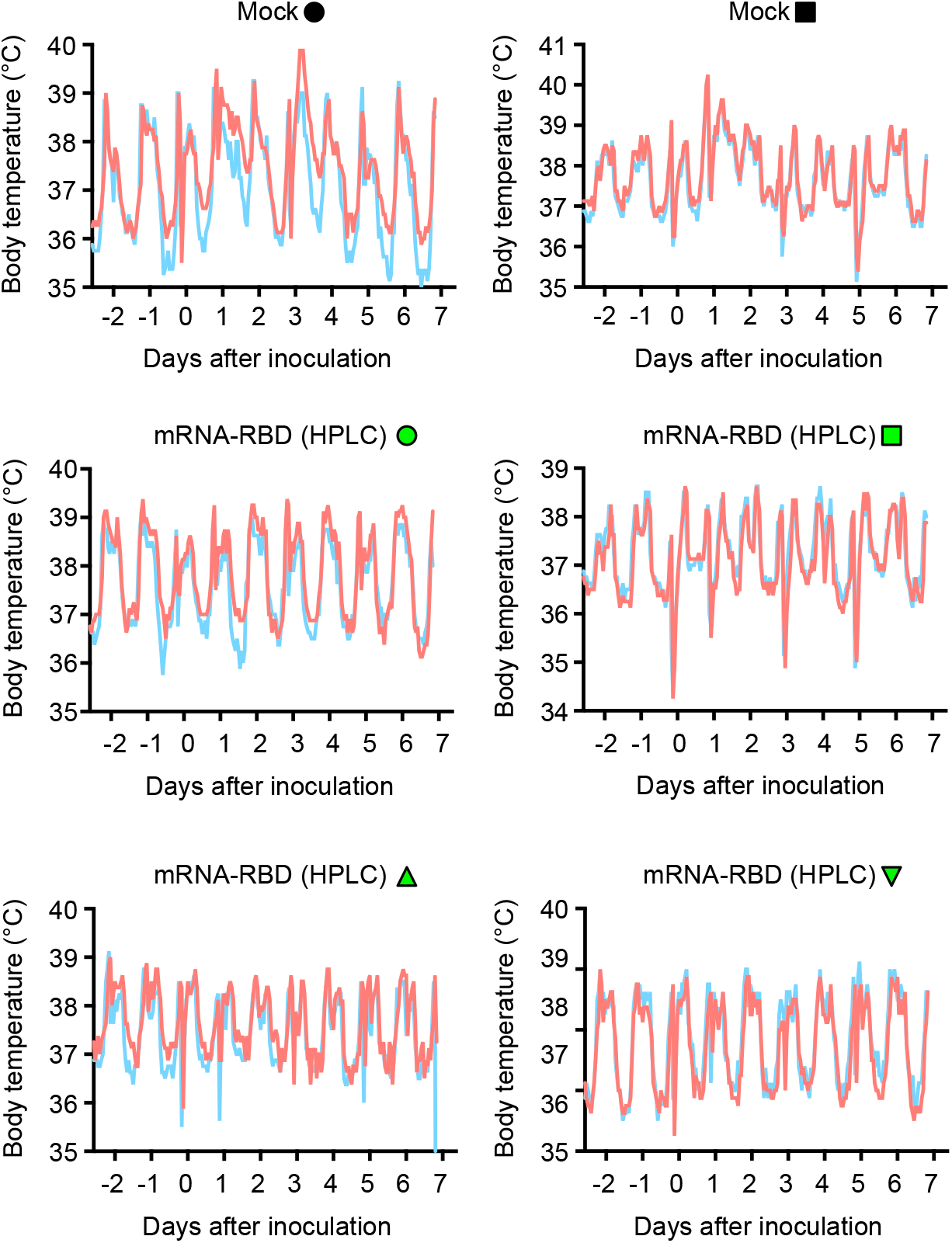
Change in body temperature after SARS-CoV-2 infection. One week after the second immunization, SARS-CoV-2 (2 × 10^7^ PFU) was inoculated into conjunctiva, nasal cavity, oral cavity, and trachea of cynomolgus. Body temperature was recorded from two days before infection using telemetry transmitters and a computer.

**Extended data Fig 11.**
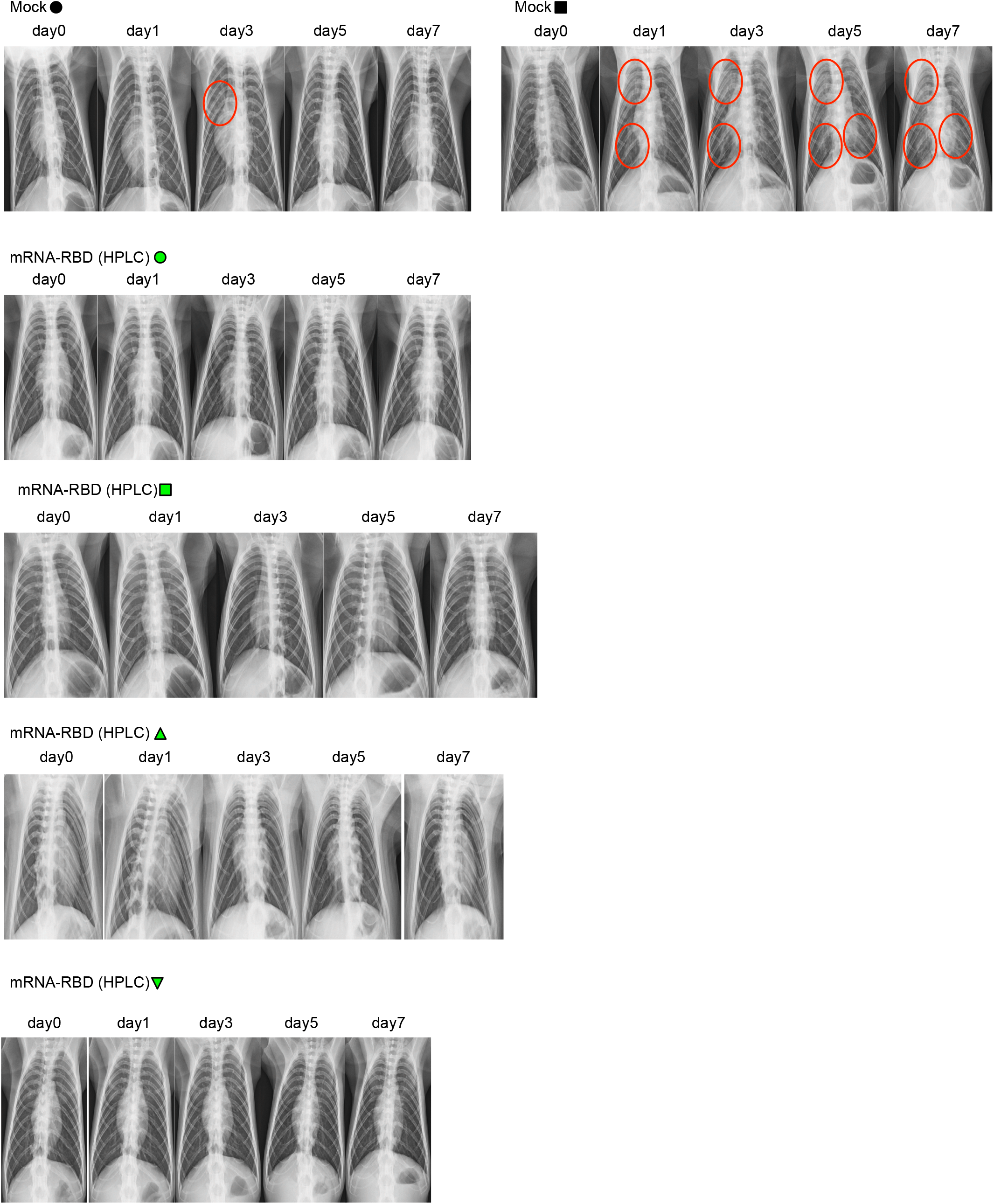
The HPLC-purified mRNA vaccine protects against SARS-CoV-2-induced pneumonia. X-ray radiographs of macaques were taken before and after SARS-CoV-2 infection.

## Material and Methods

### Mice

Six to eight week-old C57BL/6 and BALB/c mice were purchased from CLEA, Japan. The mice were maintained under specific pathogen-free conditions. All mouse studies were approved by the Animal Experiment Committee of the Institute of Medical Science, University of Tokyo.

### Cynomolgus macaque

Seven to ten-year-old female cynomolgus macaques born at Shiga University of Medical Science and originating from Philippines, Vietnam, and China were used. All procedures were performed under ketamine and xylazine anesthesia, and all efforts were made to minimize suffering. Food pellets of CMK-2 (CLEA Japan, Inc., Tokyo, Japan) were provided once a day after recovery from anesthesia and drinking water was available *ad libitum*. The animals were singly housed in cages under controlled conditions of light (12-h light/12-h dark cycle, lights on at 8:00 a.m.). The macaques were challenged with the SARS-CoV-2 (2 × 10^7^ PFU/7 mL HBSS), which was inoculated into the conjunctiva (0.05 mL × 2), nostrils (0.5 mL × 2), oral cavity (0.9 mL), and trachea (5 mL) with pipettes and catheters under ketamine/xylazine anesthesia. Under ketamine/xylazine anesthesia, two cotton sticks (Eiken Chemical, Ltd., Tokyo, Japan) were used to collect fluid samples from the conjunctivas, nasal cavities, oral cavities and tracheas, and the sticks were subsequently immersed in 1 mL of Dulbecco’s modified Eagle medium (DMEM, Nacalai Tesque, Kyoto, Japan) containing 0.1% bovine serum albumin (BSA) and antibiotics. A bronchoscope (MEV-2560; Machida Endoscope Co. Ltd., Tokyo, Japan) and cytology brushes (BC-203D-2006; Olympus Co., Tokyo, Japan) were used to obtain bronchial samples.

### LNP-mRNA vaccines

T7 RNA polymerase-mediated transcription *in vitro* was used to synthesize the mRNA from a linearized DNA template, which flanked the open-reading frames of RBD with the 5’ and 3’ untranslated regions and a poly-A tail. Messenger RNA for RBD (HPLC) was purified by reversed phase chromatography. Messenger RNA was encapsulated into lipid nanoparticles (LNP) composed of ionizable lipid, phospholipid, cholesterol, and PEG-lipid.

### Reagents

Overlapping 20-aa peptides of spike protein were synthesized and purchased from Eurofins Genomics (Ebersberg, Germany). The SARS-CoV-2 spike protein (ECD) and RBD were purchased from GenScript (Piscataway, NJ, USA).

### Virus

SARS-CoV-2 isolates were propagated in VeroE6 cells in Opti-MEM I (Invitrogen, Carlsbad, CA, USA) containing 0.3% bovine serum albumin (BSA) and 1 μg of L-1*-*tosylamide*-*2*-*phenylethyl chloromethyl ketone (TPCK)*-*treated trypsin/mL at 37°C.

### Immunization

Six to eight week-old C57BL/6 and BALB/c mice were intramuscularly immunized with mock, LNP-mRNA-RBD (3 μg), or LNP-mRNA-RBD (HPLC) (3 μg) on days 0 and 14. Two weeks after the second immunization, the popliteal lymph nodes, spleen, and blood were collected. Cynomolgus macaques were intramuscularly immunized with mock or LNP-mRNA-RBD (HPLC) (100 μg) on days 0 and 21. Blood was drawn on days 0, 7, 14, 21, and 28.

### ELISA

ECD and RBD-specific antibody titers were measured using ELISA. In brief, half-area 96-well plates were coated with ECD (1 μg/mL) or RBD (1 μg/mL) in bicarbonate buffer at 4°C. Plates were blocked with PBS containing 1% BSA for 60 min at room temperature. Plates were washed with PBST three times and incubated with diluted plasma or swab samples at room temperature for 120 min. Plates were washed with PBST three times and incubated with HRP-labeled goat anti-mouse IgG, IgG1, IgG2a, IgG2c, or mouse anti-monkey IgG at room temperature for 120 min. After washing with PBST three times, TMB substrate buffer was added, followed by incubation at room temperature for 10 min. Then, 1 N H_2_SO_4_was added to stop the reaction. OD values at 450 and 540 or 560 nm were measured using a spectrophotometer. The reciprocal value of the plasma dilution with OD_450_–OD_540_or OD_450_–OD_560_of 0.2 was defined as the antibody titer.

Single-cell suspensions of splenocytes from immunized mice were stimulated by peptide pools 1–8, ECD, and RBD protein for 24 hours. IFN-γ and IL-13 levels in the supernatant were measured using ELISA (R&D).

### GC B cell and TFH staining

Single-cell suspensions of popliteal lymph nodes were stained with LIVE/DEAD Aqua, anti-CD279 (29F.1A12), San Diego, CA, USA), anti-CD8a (53-6.7), anti-CD3e (145-2C11), anti-GL7 (GL7), anti-CD4 (RM4-5), anti-CD185 (L138D7), anti-CD38 (90), and anti-CD19 (6D5) antibodies. All antibodies were purchased from BioLegend, San Diego, CA, USA. The percentage of GC B cells and T_FH_ cells was analyzed by flow cytometry.

### Intracellular staining assay for cytokines

Single-cell suspensions of splenocytes were stimulated with peptide pools 2, 3, and 4 together with protein transport inhibitors (eBioscience, San Diego, CA, USA) for 6 h. After stimulation, the cells were stained with LIVE/DEAD Aqua for dead cells. After washing, the cells were stained with anti-CD8a (53-6.7), anti-CD4 (RM4-5: Invitrogen), anti-TCRβ (H57-597), anti-F4/80 (RM8), anti-TER-119 (TER-119), anti-CD11b (M1/70), anti-CD19 (6D5), anti-CD11c (N418), anti-NK-1.1 (PK136), and anti-CD45R/B220 (RA3-6B2) antibodies. All antibodies were purchased from BioLegend unless otherwise stated. After fixation, permeabilization by IC Fixation Buffer (eBioscience), intracellular cytokines, and CD3 were stained with anti-IFN-γ (XMG1.2), anti-IL-2 (JES6-5H4), anti-TNF-α (MP6-XT22), and anti-CD3 (17A2) antibodies. All antibodies were purchased from BioLegend. The percentage of cytokine-producing CD8^+^ and CD4^+^ T cells was determined by flow cytometry.

### Preparation and stimulation of human peripheral blood mononuclear cells

Peripheral blood mononuclear cells (PBMCs) were obtained from three SARS-CoV-2-uninfected healthy adult volunteers after obtaining informed consent. All experiments using human PBMCs were approved by the Institutional Review Board of the Institute of Medical Science, University of Tokyo. After preparation of PBMCs using Ficoll Histopaque, the cells were stimulated by LNP-mRNA-Full (0.4, 2, and 10 μg/mL), LNP-mRNA-RBD (0.4, 2, and 10 μg/mL), or LNP-mRNA-RBD (HPLC) (0.4, 2, and 10 μg/mL) for 24 h. IFN-α level in the culture supernatant was measured using ELISA (Mabtech, Stockholm, Sweden).

### Bone marrow-derived dendritic cells and stimulation

Bone marrow-derived dendritic cells (BM-DCs) were differentiated by culturing for seven days with murine GM-CSF. Cells were stimulated with LNP-mRNA-Full (0.4, 2, and 10 μg/mL), LNP-mRNA-RBD (0.4, 2, and 10 μg/mL), or LNP-mRNA-RBD (HPLC) (0.4, 2, and 10 μg/mL) for 24 h. IFN-α in the culture supernatant was measured using ELISA (Invitrogen).

### Neutralization activity against SARS-CoV-2 infection

Thirty-five microliters of virus (140 tissue culture infectious dose 50) was incubated with 35 μL of two-fold serial dilutions of sera for 1 h at room temperature, and 50 μL of the mixture was added to confluent VeroE6/TMPRSS2 cells in 96-well plates and incubated for 1 h at 37°C. After the addition of 50 μL of DMEM containing 5% FCS, the cells were further incubated for three days at 37°C. Viral cytopathic effects (CPE) were observed under an inverted microscope, and virus neutralization titers were determined as the reciprocal of the highest serum dilution that completely prevented CPE ^24^.

### Virus titration using VeroE6/TMPRSS2 for SARS-CoV-2

Confluent TMPRSS2-expressing Vero E6 cell line (JCRB Cell Bank, Japan) were incubated with diluted swab samples and 10% w/v tissue homogenate samples for 1 h. The cells were washed with HBSS and incubated with DMEM containing 0.1% BSA for three days ^25^. Virus titers were monitored using a microscope and calculated using the Reed-Muench method.

### Real-time RT-PCR for viral RNA

Viral RNA from swab samples and tissues (20 mg) was collected using a QIAamp Viral RNA Mini kit and RNeasy Mini Kit, respectively. Viral RNA was measured by real-time RT-PCR (2019-nCoV_N1-F, 2019-nCoV_N1-R, 2019-nCoV_N1-P, and TaqMan Fast Virus 1-step Master Mix) using CFX-96 (Bio-Rad, Hercules, CA, USA).

### Histological evaluation of lung section

Lungs were obtained at autopsy, and 8 lung tissue slices were collected from each macaque: one slice from each upper lobe and middle lobe and two slices from each lower lobe in bilateral lungs. They were fixed in 10% neutral buffered formalin for approximately 72 h, embedded in paraffin and cut into 3-μm-thick sections on glass slides. Sections were stained with hematoxylin and eosin (H & E) and observed under the light microscope. Histological evaluation was performed blindly by two pathologists based on a following criteria established in influenza virus infection ^26^ (0: normal lung, 1: mild destruction of bronchial epithelium, 2: mild peribronchiolar inflammation, 3: inflammation in the alveolar walls resulting in alveolar thickening, 4: mild alveolar injury accompanied by vascular injury, 5: moderate alveolar injury and vascular injury, 6, 7: severe alveolar injury with hyaline membrane-associated alveolar hemorrhage (under or over 50% of the section area)). The average score of 8 sections was calculated for each macaque, and the mean score of the two pathologists were defined as the histological score. SARS-CoV-2 N antigen was detected by a monoclonal antibody 8G8A (Bioss Inc) and secondary antibody following antigen retrieval using autoclave in pH 9 citrate buffer.

### Body temperature

Two weeks before virus inoculation, two temperature data loggers (iButton, Maxim Integrated, San Jose, CA) were implanted in the peritoneal cavity of each macaque under ketamine/xylazine anesthesia followed by isoflurane inhalation to monitor body temperature.

### X-ray radiography

Chest X-ray radiographs were taken using the I-PACS system (Konica Minolta Inc., Tokyo, Japan) and PX-20BT mini (Kenko Tokina Corporation, Tokyo, Japan).

### Statistical analysis

Statistical significance (P < 0.05) between groups was determined using the Mann–Whitney U test or ANOVA.

## Acknowledgments

We thank Fusako Ikeda, Ryoko Suda, Yoko Watanabe, Naomi Sato, Naoko Kitagawa and Hideaki Ishida for technical assistance. This study was supported by AMED under Grant Number JP20fk0108113h0001 and JP20nk0101625h0201.

## Author contribution

K.K., M.I., N.J., M.N., S.Y., Y.I., F.T., Y.K., K.J.I., designed research; K.K., M.I., N.J., M.N., K.H., K.I.H., S.Yamayoshi., J.T., M.I., S.Yamada., T.W., M.K., H.N., H.I., Y.K., C.T.N., Y.I., performed research; K.K., M.I., M.N., K.H., J.T., B.T., C.T.N., Y.I., analyzed data; N.J., T.N., T.S., F.T., contributed to provide LNP-mRNA vaccine. K.K., N.J., Y.I., Y.K., F.T., K.J.I. wrote the paper.

